# Novel quinolone resistance determinant, *qepA8*, in *Shigella flexneri* isolated in the United States, 2016

**DOI:** 10.1101/726950

**Authors:** Hattie E. Webb, Kaitlin A. Tagg, Jessica C. Chen, Justin Kim, Rebecca Lindsey, Louise K. Francois Watkins, Beth E. Karp, Yo Sugawara, Jason P. Folster

## Abstract

A *qepA8*+ *Shigella flexneri* was cultured from the stool of a traveler returning from India and East Asia. This chromosomally encoded *qepA* variant, has a six-base insertion, and may have been mobilized as part of a complex IS*1-*mediated composite transposon including *catA1, aadA1*, and *bla*_OXA-1_. In laboratory *E. coli*, *qepA8* alone only conferred decreased ciprofloxacin susceptibility; however, it may work in combination with additional mechanisms to confer clinical resistance.

---

Shigellosis is an important public health problem. *Shigella* causes an estimated 500,000 cases of diarrhea in the United States annually (1). In both the United States and Europe, children in daycare centers, travelers to developing countries, and men who have sex with men are most often infected. Antibiotics are used to treat severe infection and to shorten the duration of illness (2). Although not usually necessary, ciprofloxacin (for adults) and azithromycin (for children) are the preferred treatment (3, 4). Increasing resistance poses a major therapeutic challenge (5). In Enterobacteriaceae, quinolone resistance is largely attributed to mutations in the quinolone resistance-determining regions (QRDR) of *gyrA*, *gyrB*, *parC*, and *parE*, and plasmid-mediated quinolone resistance (PMQR) determinants (e.g., *qnr* genes, *aac(6’)-Ib-cr*, or *qepA*). The *qep*A gene encodes a 14-transmembrane-segment efflux pump of the major facilitator superfamily. The functionality of four variants, designated *qepA1-4*, have been described in literature (6–10); however, other *qepA* sequence variations were recently designated *qepA5-10*, despite having unknown functionality against quinolones (11). Here, we describe a *qepA8* detected in *Shigella flexneri* from a patient in the United States.

In the United States, public health laboratories submit every twentieth *Shigella* isolate to the Centers for Disease Control and Prevention (CDC) National Antimicrobial Resistance Monitoring System (12). In 2016, 544 *Shigella* isolates were genome sequenced and phenotypically tested against 14 antibiotics by broth microdilution (Sensititre™; Table 1). During routine screening, a single ciprofloxacin-resistant *Shigella flexneri* strain had a *qepA* variant with a six base insertion at position 1,065. The *S*. *flexneri* was cultured from the stool of a 69-year-old man in 2016. The patient reported spending several months traveling in Asia (Taiwan, Thailand, Laos, Cambodia, and India) and sought treatment at a local hospital in India after developing abdominal cramping and bloody diarrhea. According to the patient, diarrheagenic *Escherichia coli* was cultured from his stool and was prescribed ciprofloxacin in India, but his diarrhea worsened and he sought medical care in the United States 2-3 weeks later. However, in the United States, *Shigella* was isolated from his stool and a five-day course of ciprofloxacin was again prescribed. The patient was not hospitalized and recovered. Due to the close genetic relationship between *E. coli* and *Shigella*, it is possible the patient had a single prolonged infection of *Shigella*, both ciprofloxacin treatments were ineffective, and the *Shigella* infection resolved on its own after >4 weeks. Genomic DNA was extracted using the Wizard Genomic DNA Purification (Promega Corporation, Madison, WI, USA). Libraries sequenced on the Illumina HiSeq platform were prepared using the NEBNext Ultra DNA library prep kit (New England Biolabs, Ipswich, MA). Libraries sequenced on the Pacific Biosciences RSII system were generated using standard library protocols of the DNA template preparation kit (PacBio, Menlo Park, CA, USA). Sequence reads were filtered and assembled *de novo* using the Hierarchical Genome Assembly Process (HGAP) v3 (13) and circularized using Circlator v1.5.5 (14). PacBio sequences were corrected using Illumina reads. Annotation was performed using Prokka (v1.13.3 (15)) and Galileo AMR (16) before submission to NCBI (2016AM-0877; Accession no. CP033510.1 and CP033511.1). Resistance genes were identified using assembly-based methods with the ResFinder database (https://github.com/phac-nml/staramr) and QRDR mutations by direct sequence comparison.

**TABLE 1.**
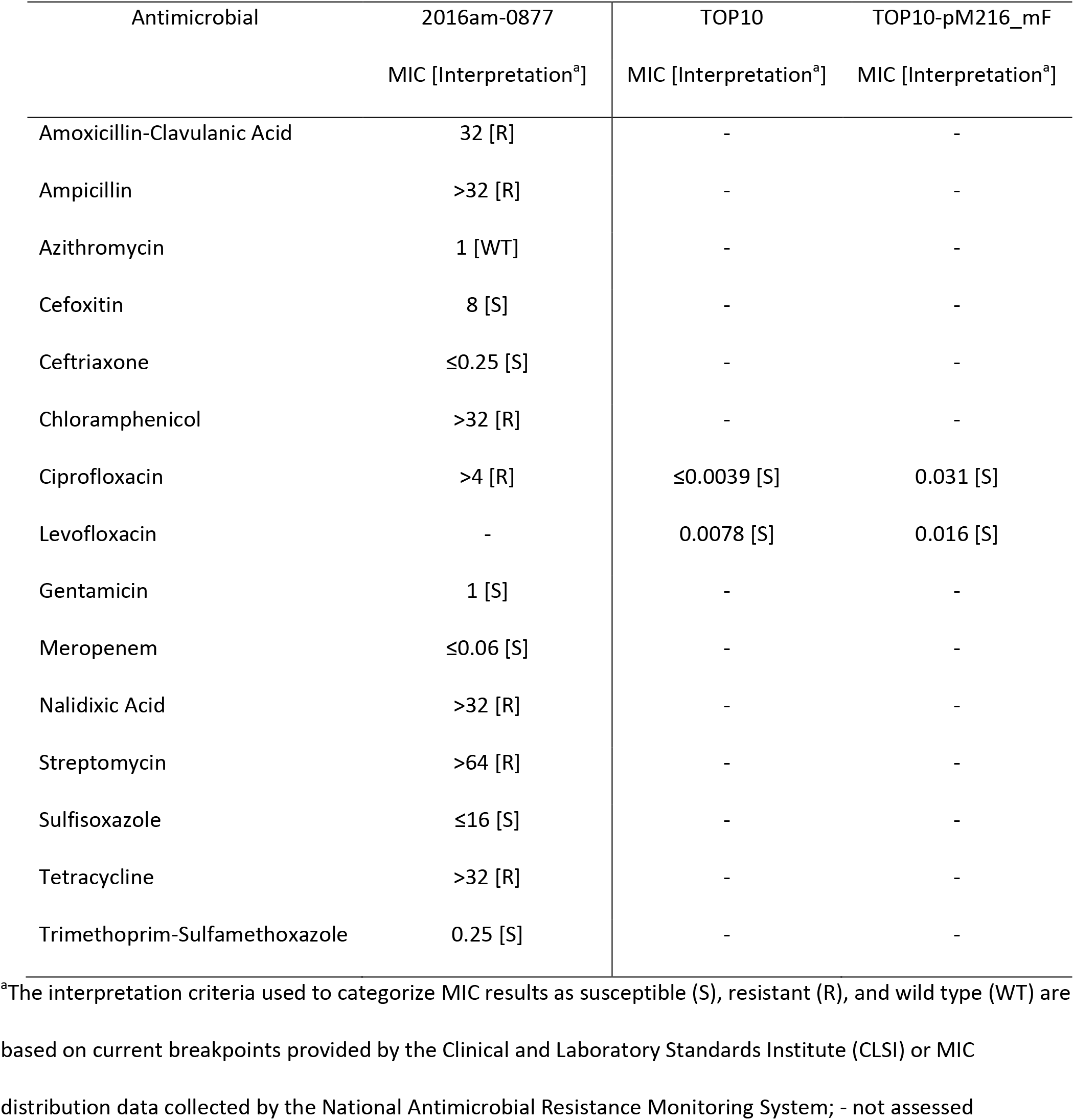
Minimum Inhibitory Concentrations (MIC; µg/mL) and interpretations for tested antimicrobials against 2016AM-0877, *E. coli* TOP10, and transformant TOP10-pM216_mF (in triplicate)

In 2016AM-0877, *qepA8* was chromosomally located and may have been captured/mobilized as part of an IS*1*-mediated composite transposon flanked by direct repeats (Figure 1), which also includes *catA1* (V00622), *aadA1* (JX185132), *bla*_OXA-1_ (J02967), and a truncated *dfrB4* gene. In addition, QRDR mutations were observed in *gyrA* (Ser83Leu and Asp87Asn) and *parC* (Ser80Ile). Additional resistance genes (outside the *qepA*-transposon) were also identified, including two *aadA1* (JQ480156; JX185132), *dfrA1* (X00926), *bla*_OXA-1_ (J02967), and two *tet*(B) (AF326777). Phenotypic resistance was observed to amoxicillin-clavulanic acid, ampicillin, chloramphenicol, ciprofloxacin, nalidixic acid, streptomycin, and tetracycline (Table 1).

**Figure 1.**
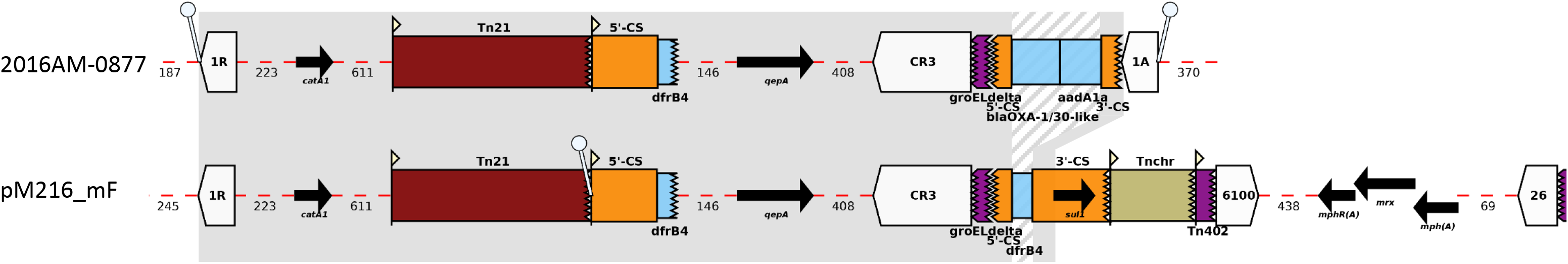
Comparative analysis of the genetic context of *qepA* variant in the chromosome of 2016AM-0877 (CP033510.1, reported herein) and on pM216_mF (18). Gray indicates regions shared between both strains and striped regions indicates differing cassette arrays, where variability may be expected. Gene features are shown as arrows pointed in the orientation of the features (*catA1*, *qepA*, *sul1*, *mphR*(A)/(A), *mrx*). White block arrows represent insertion sequences (IS*1R/A*, IS*CR3*, IS*6100*, IS*26*), orange boxes represent conserved segments of integrons (5′-CS and 3′-CS), light blue boxes represent gene cassettes (*bla*_OXA-1_, *aadA1*, *dfrB4*), and ‘lollipops’ represent direct repeats flanking each end of the inserted region. Unit transposons (Tn*21*, Tn*chr*, Tn*402*) are shown as colored boxes and their inverted repeat regions are shown as flags, with the flat side as the outer boundary of the transposon. Truncations are represented by a jagged edge on the truncated side(s) of the feature and the number of bases present in between features is shown beneath the red dashed line. Diagram modified from that generated by Galileo AMR (16).

A NCBI Pathogen Detection Browser query (https://www.ncbi.nlm.nih.gov/pathogens; May 2019) yielded 24 *qepA8*-positive *E. coli* and *Shigella*. Although, 2016AM-0877 was the first observed by Pathogen Detection (create date November 11, 2018), isolates cultured from specimens collected as early as 2010 were uploaded to NCBI and later found to be *qepA8*-positive. A majority of the *qepA8*-positive strains (n=15) were of clinical or environmental origin and were collected in Myanmar during August 2015-January 2017 (17, 18). Of the publicly available long-reads, one had a 149,768-bp plasmid—herein referred to as pM216_mF (Genbank LC492469)— which carried *qepA8* in a complex composite transposon that appeared to be highly related (<10 single-nucleotide polymorphisms in the shared region [gray in Figure 1]) to that observed in 2016AM-0877. The similarities suggest that a pM216_mF-like plasmid was likely the source of the context observed in the chromosome of 2016AM-0877, which may have initially been mobilized by IS*1*.

Since *qepA8* was chromosomally-encoded in 2016AM-0877 alongside QRDR mutations, a transformation experiment was used to assess functionality of plasmid-encoded *qepA8* in M216. Briefly, plasmid DNA was extracted from M216 and introduced into a TOP10 *E. coli* strain by electroporation. Transformants were selected on LB plates containing 30 μg/mL of chloramphenicol. Plasmid size and *qepA* presence was confirmed by S1-PFGE (17) and PCR (5’-ATGTCCGCCACGCTCCACGAC-3’, 5’-TCAACCAGATGCGAGCGCTG-3’), respectively. Broth microdilution for the TOP10 transformants compared with untransformed E. coli TOP10 (performed in triplicate) found an at least eight-fold and a two-fold increase in MIC was observed for ciprofloxacin and levofloxacin (Table 1), respectively; however, MIC values remained significantly below clinical breakpoints (19).

In conclusion, we describe a novel *qepA* variant that mediates reduced susceptibility to ciprofloxacin. Although the ciprofloxacin MIC did not increase above the resistance breakpoint, PMQR genes may complement other quinolone resistance mechanisms to reach clinical resistance levels, and may additionally facilitate the selection of higher-level resistance by increasing the mutant prevention concentration (20). To our knowledge, this is the first report on the genetic context and functionality of *qepA8*. Publicly available sequences indicate *qepA8* has also been reported in *E. coli*/*Shigella* outside of the United States and was present as early as 2010. As public health labs transition into this era of whole genome sequencing, there will be increasing opportunities to investigate novel variants of resistance genes and determine the extent to which they confer antimicrobial resistance.

## Acknowledgments

We thank state and local health departments for collecting patient information and submitting isolates. The findings and conclusions in this report are those of the authors and do not necessarily represent the official position of the CDC.

## References

1. Scallan E, Hoekstra RM, Angulo FJ, Tauxe RV, Widdowson MA, Roy SL, Jones JL, Griffin PM. 2011. Foodborne illness acquired in the United States-major pathogens. Emerging Infect Dis 17:7–15.

2. Niyogi SK. 2005. Shigellosis. The journal of microbiology 43:133–143.

3. Shane AL, Mody RK, Crump JA, Tarr PI, Steiner TS, Kotloff K, Langley JM, Wanke C, Warren CA, Cheng ACJCID. 2017. 2017 Infectious Diseases Society of America clinical practice guidelines for the diagnosis and management of infectious diarrhea. 65:e45–e80.

4. American Academy of Pediatrics. 2018. Section 3: Summaries of Infectious Diseases, p 723–727. In Kimberlin D, Brady M, Jackson M, Long S (ed), Red Book: 2018 Report of the Committee on Infectious Diseases, 31st ed. American Academy of Pediatrics, Itasca, IL.

5. Christopher PRH, David KV, John SM, Sankarapandian V. 2010. Antibiotic therapy for *Shigella* dysentery. Cochrane Database of Systematic Reviews doi: 10.1002/14651858.CD006784.pub4.

6. Périchon B, Courvalin P, Galimand M. 2007. Transferable resistance to aminoglycosides by methylation of G1405 in 16S rRNA and to hydrophilic fluoroquinolones by QepA-mediated efflux in *Escherichia coli*. Antimicrobial agents and chemotherapy 51:2464–2469.

7. Yamane K, Wachino J-i, Suzuki S, Kimura K, Shibata N, Kato H, Shibayama K, Konda T, Arakawa Y. 2007. New plasmid-mediated fluoroquinolone efflux pump, QepA, found in an *Escherichia coli* clinical isolate. Antimicrobial agents and chemotherapy 51:3354–3360.

8. Wang D, Huang X, Chen J, Mou Y, Qi Y. 2017. Characterization of a novel *qepA3* variant in *Enterobacter aerogenes*. J Microbiol Immunol Infect 50:254–257.

9. Cattoir V, Poirel L, Nordmann P. 2008. Plasmid-mediated quinolone resistance pump QepA2 in an *Escherichia coli* isolate from France. Antimicrobial agents and chemotherapy 52:3801–3804.

10. Manageiro V, Félix D, Jones-Dias D, Sampaio DA, Vieira L, Sancho L, Ferreira E, Caniça M. 2017. Genetic Background and Expression of the New *qepA4* Gene Variant Recovered in Clinical TEM-1-and CMY-2-Producing *Escherichia coli*. Frontiers in microbiology 8:1899.

11. Ruiz JJREdQ. 2018. In silico analysis of transferable QepA variants and related chromosomal efflux pumps. 31:537.

12. Centers for Disease Control and Prevention (CDC). 2018. National Antimicrobial Resistance Monitoring System for Enteric Bacteria (NARMS): Human Isolates Surveillance Report for 2015 (Final Report). U.S. Department of Health and Human Services, CDC, Atlanta, Georgia.

13. Chin C-S, Alexander DH, Marks P, Klammer AA, Drake J, Heiner C, Clum A, Copeland A, Huddleston J, Eichler EEJNm. 2013. Nonhybrid, finished microbial genome assemblies from long-read SMRT sequencing data. 10:563.

14. Hunt M, De Silva N, Otto TD, Parkhill J, Keane JA, Harris SRJGb. 2015. Circlator: automated circularization of genome assemblies using long sequencing reads. 16:294.

15. Seemann T. 2014. Prokka: rapid prokaryotic genome annotation. Bioinformatics 30:2068–2069.

16. Partridge SR, Tsafnat GJ. 2018. Automated annotation of mobile antibiotic resistance in Gram-negative bacteria: the Multiple Antibiotic Resistance Annotator (MARA) and database. Journal of Antimicrobial Chemotherapy 73:883–890.

17. Sugawara Y, Akeda Y, Hagiya H, Sakamoto N, Takeuchi D, Shanmugakani RK, Motooka D, Nishi I, Zin KN, Aye MM. 2019. Spreading patterns of NDM-producing Enterobacteriaceae in clinical and environmental settings in Yangon, Myanmar. J Antimicrobial agents 63:e01924–18.

18. Sugawara Y, Akeda Y, Sakamoto N, Takeuchi D, Motooka D, Nakamura S, Hagiya H, Yamamoto N, Nishi I, Yoshida H. 2017. Genetic characterization of *blaNDM*-harboring plasmids in carbapenem-resistant *Escherichia coli* from Myanmar. PloS one 12:e0184720.

19. Clinical and Laboratory Standards Institute. 2012. CLSI document M07-A9: Methods for Dilution Antimicrobial Susceptibility Tests for Bacteria That Grow Aerobically; Approved Standard, 9th ed. Clinical and Laboratory Standards Institute, Wayne, PA.

20. Rodríguez-Martínez JM, Machuca J, Cano ME, Calvo J, Martinez-Martinez L, Pascual A. 2016. Plasmid-mediated quinolone resistance: two decades on. Drug Resistance Updates 29:13–29.

